# TeloSearchLR: an algorithm to detect novel telomere repeat motifs using long sequencing reads

**DOI:** 10.1101/2024.10.29.617943

**Authors:** George Chung, Fabio Piano, Kristin C Gunsalus

## Abstract

Telomeres are eukaryotic chromosome end structures that guard against sequence loss and aberrant chromosome fusions. Telomeric repeat motifs (TRMs), the minimal repeating unit of a telomere, vary from species to species, with some evolutionary clades experiencing a rapid sequence divergence. To explore the full scope of this evolutionary divergence, many bioinformatic tools have been developed to infer novel TRMs using repetitive sequence search on short sequencing reads. However, novel telomeric motifs remain unidentified in up to half of the sequencing libraries assayed with these tools. A possible reason may be that short reads, derived from extensively sheared DNA, preserve little to no positional context of the repetitive sequences assayed. On the other hand, if a sequencing read is sufficiently long, telomeric sequences must appear at either end rather than in the middle. The TeloSearchLR algorithm relies on this to help identify novel TRMs on long reads, in many cases where short-read search tools have failed. In addition, we demonstrate that TeloSearchLR can reveal unusually long telomeric motifs not maintained by telomerase, and it can also be used to anchor terminal scaffolds in new genome assemblies.

## INTRODUCTION

Replicative DNA polymerases lack the ability to synthesize the extreme 5’ end of double-stranded linear DNA. Because of this, repeated cycles of DNA replication would lead to a progressive loss of chromosome ends, a phenomenon known as the End Replication Problem^1,2^. Several solutions to this problem exist in nature, including covalently closed hairpin ends found in certain bacteria and phages^3–9^, and DNA synthesis primed by covalently linked terminal proteins found in several virus families^10–13^. In eukaryotes, the most common solution is the regulated addition of tandemly repeated sequences to the 3’ end by telomerase^14^, circumventing the End Replication Problem by allowing more of the complementary 5’ end to be replicated.

Telomerase is a protein-RNA complex that synthesizes telomeric repeats using a short template on the telomerase RNA^15^. Even though the telomerase RNAs across the tree of life are not particularly well conserved in size or in primary sequence, the template region – and thus the telomeric repeat motif (TRM) - appears surprisingly constant^16,17^: TTTAGGG in most flowering plants^18–20^ and TTAGGG in most metazoans^21,22^ (except insects where the motif is mostly TTAGG^23–25^). Nevertheless, certain clades appear to experience an accelerated divergence in their TRMs. Species in the genus *Allium* (onions, garlic, and relatives) have a CTCGGTTATGGG motif^18,26^. Similarly, TRMs in coleopteran (beetles)^23,24,27,28^ and hymenopteran^29^ (sawflies, bees, wasps and ants) species have repeatedly diverged from the ancestral insect motif.

Determining the full diversity of TRMs can allow us to find specific molecular drivers behind this diversity. As such, many different strategies have been deployed to detect telomeric repeats across species – mostly based on hybridization assays or bioinformatic analyses. Hybridization-based assays use labelled DNA probes to anneal to telomeric sequences: being the gold standard for verifying known telomeric motifs (especially in combination with the Bal-31 exonuclease^30^), these assays are not suitable for identifying new and divergent telomeric repeat motifs^28^, because generating probes for every possible sequence permutation that can appear at telomeres is impractical. Thus, bioinformatic strategies have been more successful at identifying novel TRMs.

Because most telomeres maintained by telomerase are tandem repeat clusters of short (5∼30 bp) motifs, most bioinformatic strategies rely on tandem repeat detection, using Tandem Repeats Finder^31^, mreps^32^, the Telomeric Repeats Identification Pipeline^29^ or a similar algorithm on short sequencing reads generated on the Illumina sequencing platform. These strategies rank the tandem repeats by their occurrence in the sequencing library. In species with relatively long telomeres, the telomeric motif is often one of the most frequent tandem repeats. Indeed, this strategy has successfully identified novel telomeric repeat motifs in many species of fungi^33,34^, plants^35^, insects^28,29^, and nematodes^36^.

These short-read search strategies are not without limitations. When TRMs are long – such as the one from *Kluyveromyces lactis* (25-bp long) – short read searches fail because too few repeats are found on short sequencing reads^34^. Furthermore, because these search strategies look for repeat motifs with the most frequent occurrences on sequencing reads, extremely short telomeres with very low repeat copy numbers may also be missed. Perhaps due to these and other limitations, a search strategy using Tandem Repeats Finder could not detect the telomeric repeat motif in 23 out of 96 Coleopteran species surveyed^28^, and a similar search strategy using the Telomeric Repeats Identification Pipeline failed to detect the telomeric repeat motif in 66 out of 129 insect species surveyed^29^.

To fill in the gaps in the survey of telomeric motif diversity, we developed TeloSearchLR (<u>Telo</u>mere <u>Search</u> on <u>L</u>ong <u>R</u>eads), a new telomeric repeat motif search strategy that circumvents the limitations of the older search strategies by taking advantage of a growing number of publicly available long-read genomic sequencing libraries. We successfully used the algorithm to identify telomere repeat motifs where they were previously unknown. In this report, we also describe when our search strategy failed – likely due to telomeres being much longer than the sequencing reads. We further propose that TeloSearchLR can be used to identify subtelomeric repeat sequences present on long sequencing reads.

## METHODS AND MATERIALS

### Algorithm design and key parameters

TeloSearchLR is a Python script that surveys long genomic sequencing reads for telomeric repeats. Long sequencing reads generated by either the Pacific Biosciences platform (PacBio) or the Oxford Nanopore Technologies platform (ONT) capture longer repeating motifs than short sequencing reads. Moreover, the lack of extensive DNA fragmentation in long-read sequencing preserves natural DNA ends – such as replication intermediates, DNA repair intermediates, or telomeres. Intact telomeres always appear in their canonical orientation (usually the “G-rich strand”) at the 3’ ends of long sequencing reads, and as their reverse complement (usually the “C-rich strand”) at the 5’ ends - a phenomenon we call “terminal stranded occupancy”. Using this logic, the TeloSearchLR algorithm helps us search for bona fide telomere repeat motifs (TRMs) by plotting the motif occupancy patterns at the 5’ and the 3’ ends of sequencing reads.

TeloSearchLR involves two major steps: ranking and plotting. In the ranking step, TeloSearchLR uses TideHunter (v1.5.4)^37^ to identify all tandem repeat motifs appearing within the first and the last *t* bp on all reads 2*t* bps or longer (default *t* = 1000, specified by the -t flag, **Fig. 1a**). The user specifies the range of repeat periods (in bps) for the algorithm to consider, using the -k (shortest period) and the -K (longest period) flags. TideHunter then produces a tabular output detailing the detected repeat motifs and their locations on the sequencing reads. Using this tabular output, TeloSearchLR generates a list of motifs ranked from the frequent to the least frequent (**Fig. 1a**).

**Figure 1:**
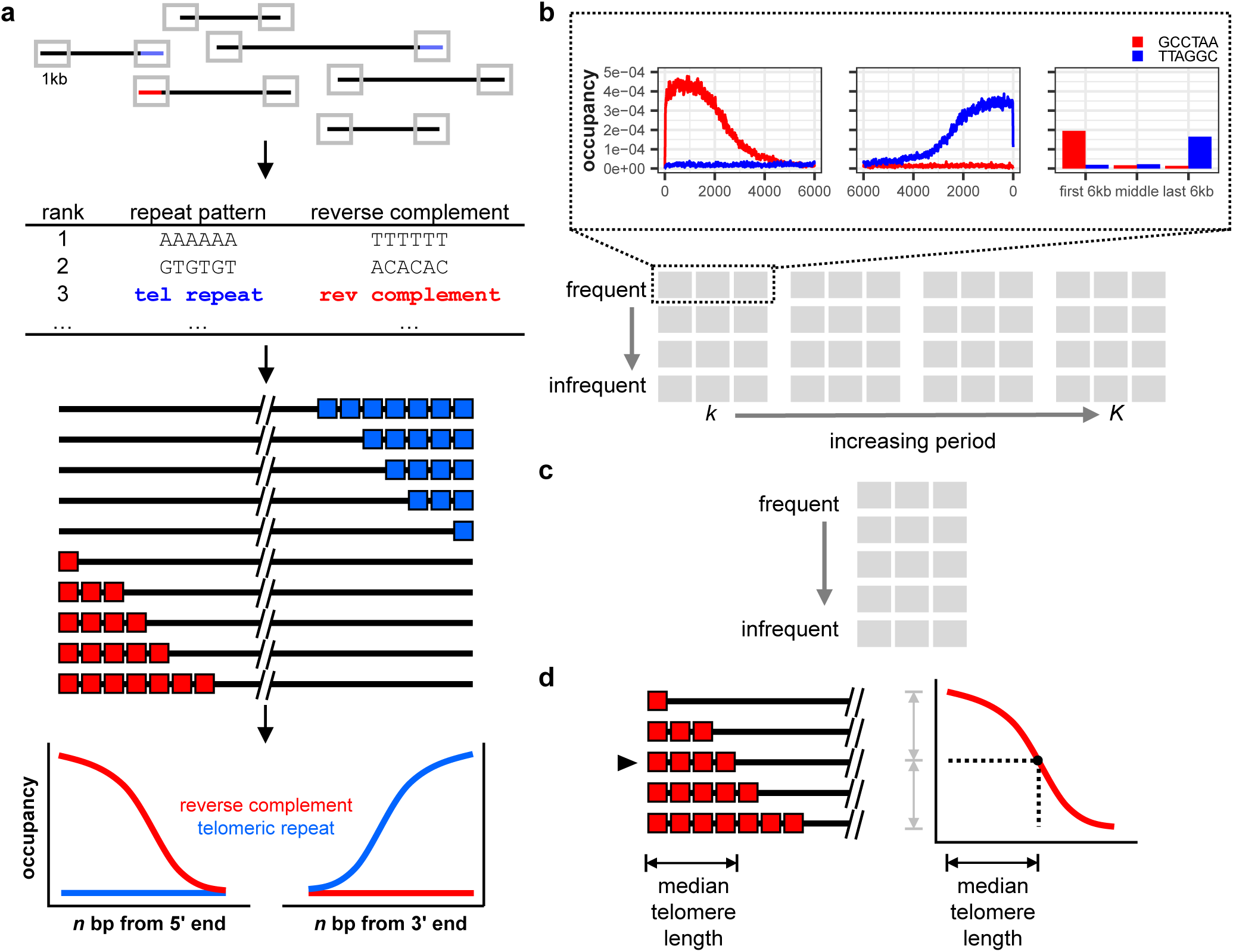
Algorithm design and the expected output. **a,** The TeloSearchLR algorithm examines the terminal 1 kb of long-read sequences for tandem repeats and ranks the repeat motif classes by how frequently they appear in a sequencing library. As natural dsDNA ends, telomeric repeat motifs (TRMs) are expected to be among the most abundant repeat classes and should show a strand bias: the conventional motif orientation (pointing toward the chromosome end) should be enriched at the 3’ ends of sequencing reads, while the reverse complement should be enriched at the 5’ ends. Visual inspection of the terminal stranded occupancy pattern should distinguish TRMs from other repeat motifs. **b,** Algorithm output in ‘exhaustive’ mode. Occupancy patterns are graphed by distance from the 5’ and 3’ ends of sequencing reads and by terminal vs. middle regions. Patterns are arranged by repeat motif period in columns, and by total occupancy in rows. **c,** Algorithm output in ‘occupancy’ mode, which ranks repeat motifs by frequency irrespective of their period. **d,** The occupancy pattern naturally reflects the distribution of telomere lengths, such that the *x* value (bp) at half the maximum *y* value (occupancy) should be the median telomere length. Thus, telomere lengths may be estimated from the algorithm output.

In the plotting step, TeloSearchLR iterates through the list of ranked motifs and determines the positions where each motif and its reverse complement are occupying on the sequencing reads, this time using the tabular output of a TideHunter search across entire reads (rather than just the terminal *t* bps). TeloSearchLR will plot the occupancy patterns for the most frequent motif (numerical ranking specified by the -m flag) to the least (-M flag followed by a number). The user also specifies a plotting window using the -n flag so that TeloSearchLR records the positions occupied by repeat motifs in the first and last *n* nucleotides as well as the middle of all sequencing reads 2*n* bp or longer. At the end of the plotting step, the putative TRMs can then be identified visually by their terminal stranded occupancy – where the reverse complement of the TRM is almost exclusively found at the first *n* bp of the reads, and the TRM is almost exclusively found at the last *n* bp of the reads (**Fig. 1a**). As read lengths are variable, the region excluding the first and last *n* bp is also variable. Thus, the occupancy fraction of a repeat motif in the middle of the reads is represented in a separate bar graph, and a bona fide TRM should be underrepresented in the middle of sequencing reads (**Fig. 1b**).

TeloSearchLR can output the plots by the repeat motif period *and* by the motif occupancy (the “exhaustive mode”, enabled by the -e flag, **Fig. 1b**) or by occupancy only (the “occupancy mode”, by default, **Fig. 1c**). Sorting the plots by repeat period is useful when searching for repeat motifs of a known length, while sorting by occupancy is useful when the telomeric repeat period is unknown. Because the TRM occupancy plot reflects the distribution of telomere lengths, we can estimate the median telomere length by determining the *x* value at which the corresponding occupancy value (*y*) is roughly one-half of the maximum repeat occupancy on the same plot (**Figure 1d**). The run commands for every library examined in this report are provided in **Table 1**, and the TRMs found by our algorithm, as well as the estimated median telomere lengths for various species, are reported in **Table 2**.

**Table 1:**
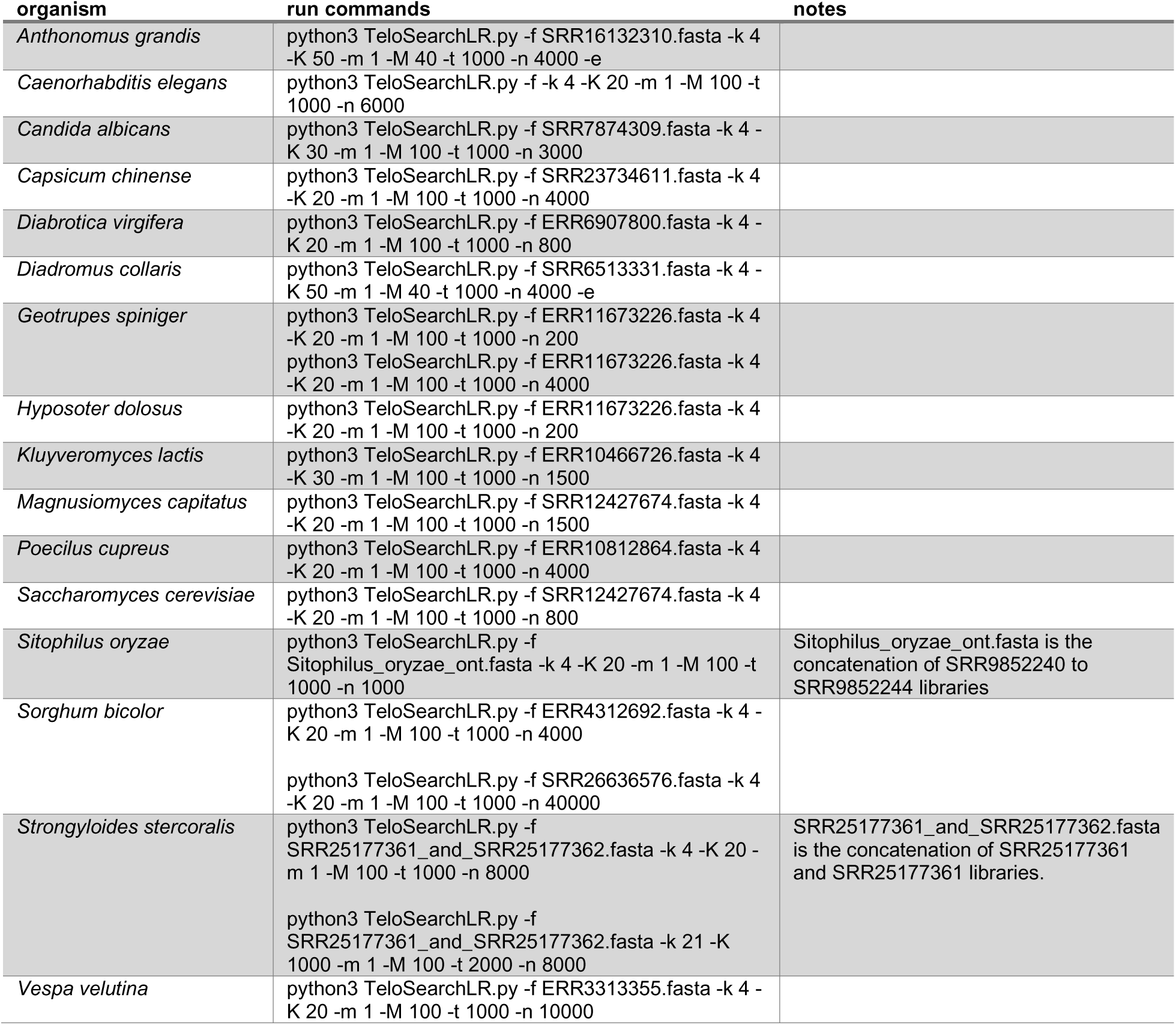
Commands for TeloSearchLR analysis

**Table 2:**
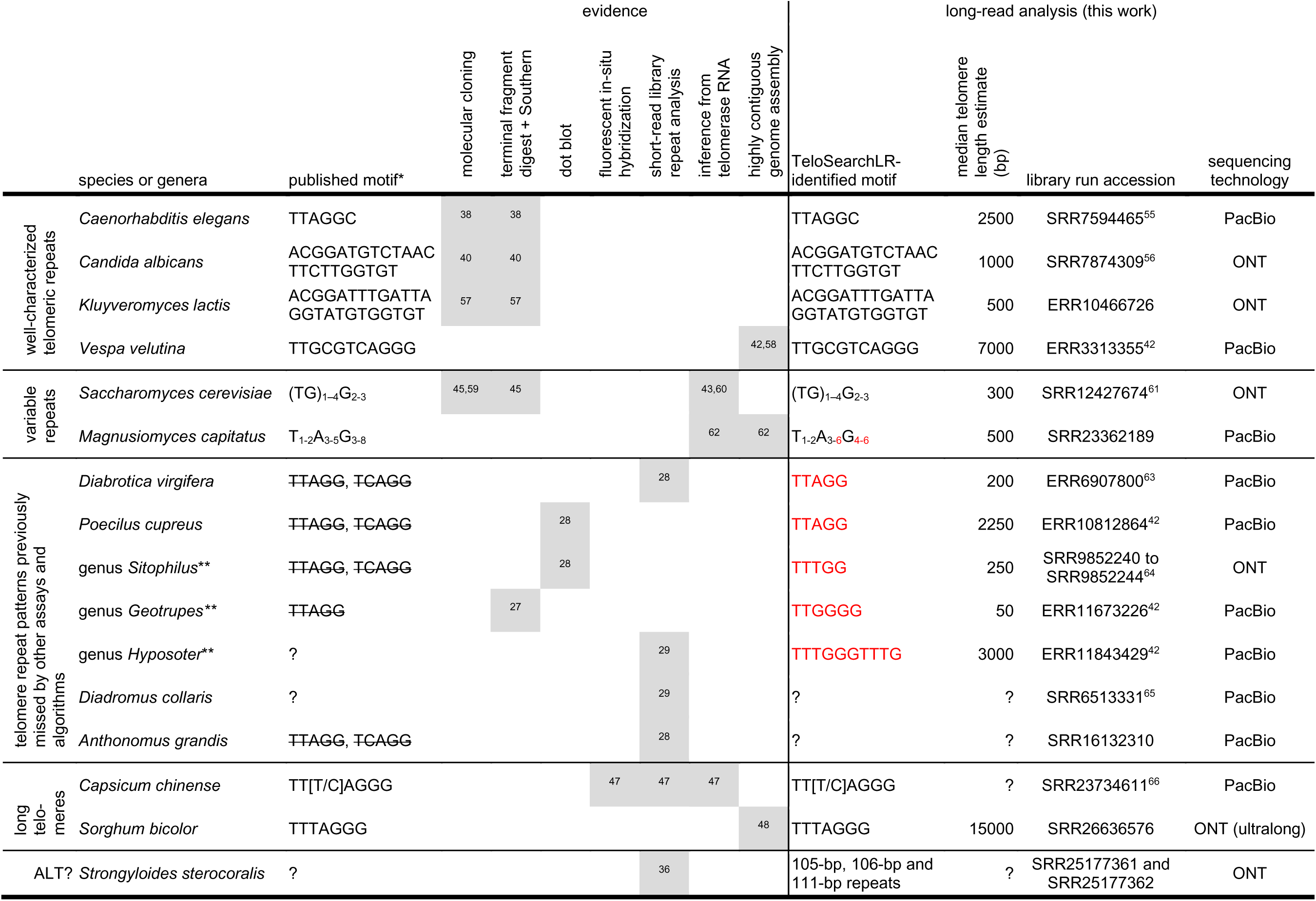
Telomeric repeat motifs identified visually from terminal stranded occupancy compared to telomeric repeats identified by molecular and older bioinformatic methods. Patterns in red are novel candidate motifs identified using TeloSearch-LR. *patterns with a strikethrough (indicate repeat motifs that have been ruled out. **Previously tested species were *Hyposoter didymator, Sitophilus granrius,* and *Geotrupes stercorarius*. Species tested for this work are *Hyposoter dolosus*, *Sitophilus oryzae*, and *Geotrupes spiniger*.

We provide one additional way of running TeloSearchLR we named “single-motif mode”, enabled by the - s flag followed by the repeat motif. In this mode, TeloSearchLR only graphs the occupancy of a single repeat motif supplied by the user, thus skipping the ranking step altogether. This is useful for quickly checking if a specific motif is telomeric. Required parameters are the fasta read file (-f), the sequence motif to be examined (-s), the graphing window (-n), and a TideHunter tabular output file (-T), which can be from an earlier run of TeloSearchLR, or from a TideHunter run independent from TeloSearchLR.

### Sequencing libraries

Sequencing libraries were downloaded from the NCBI Sequence Read Archive (https://www.ncbi.nlm.nih.gov/sra) using the fasterq-dump command from SRAtoolkit (v3.0.5, https://github.com/ncbi/sra-tools/). Since sequencing quality information is not needed for TeloSearchLR, the libraries were downloaded in FASTA format using the --fasta option. **Table 2** lists the accession numbers and the relevant references for these libraries.

## RESULTS

### Detection of well-characterized telomeric repeat motifs

We benchmarked our algorithm on long genomic sequencing reads from organisms whose TRMs were previously identified by different methods. We aimed to demonstrate that our search strategy could detect a range of telomeric repeat motifs from four organisms – from 6-mer motifs (the nematode *Caenorhabditis elegans*^38^) to 25-mer motifs (the yeast *Kluyveromyces lactis*^39^) (**Table 2**). The telomeric repeat motif of three species (*C. elegans*, *K. lactis*, and *C. albicans*) were initially discovered through the molecular cloning of chromosome ends and verified with terminal restriction fragment digests^38–40^. The motif for the wasp *Vespa velutina* was discovered through an examination^41^ of a highly contiguous genome assembly generated by the Darwin Tree of Life project^42^. In all occupancy mode runs on *C. elegans*, *C. albicans, K. lactis,* and *V. velutina* long sequencing reads, the most frequent repeat motif with a clearly terminal stranded occupancy pattern is the previously identified TRM with its reverse complement, confirming that TeloSearchLR can help us identify the telomeric repeat motif (**Table 2**, **Fig. 2a-d**, **Supplementary fig. 1-4**). Several other repeat motifs also show a terminal and stranded occupancy pattern (**Supplementary fig. 1-4**, pink boxes). These repeat motifs have a Hamming distance of 1 or 2 to the known telomeric repeat motif and appear at a much lower frequency: thus, these motifs likely represent sequencing errors or mutations to the telomeric sequences.

**Figure 2:**
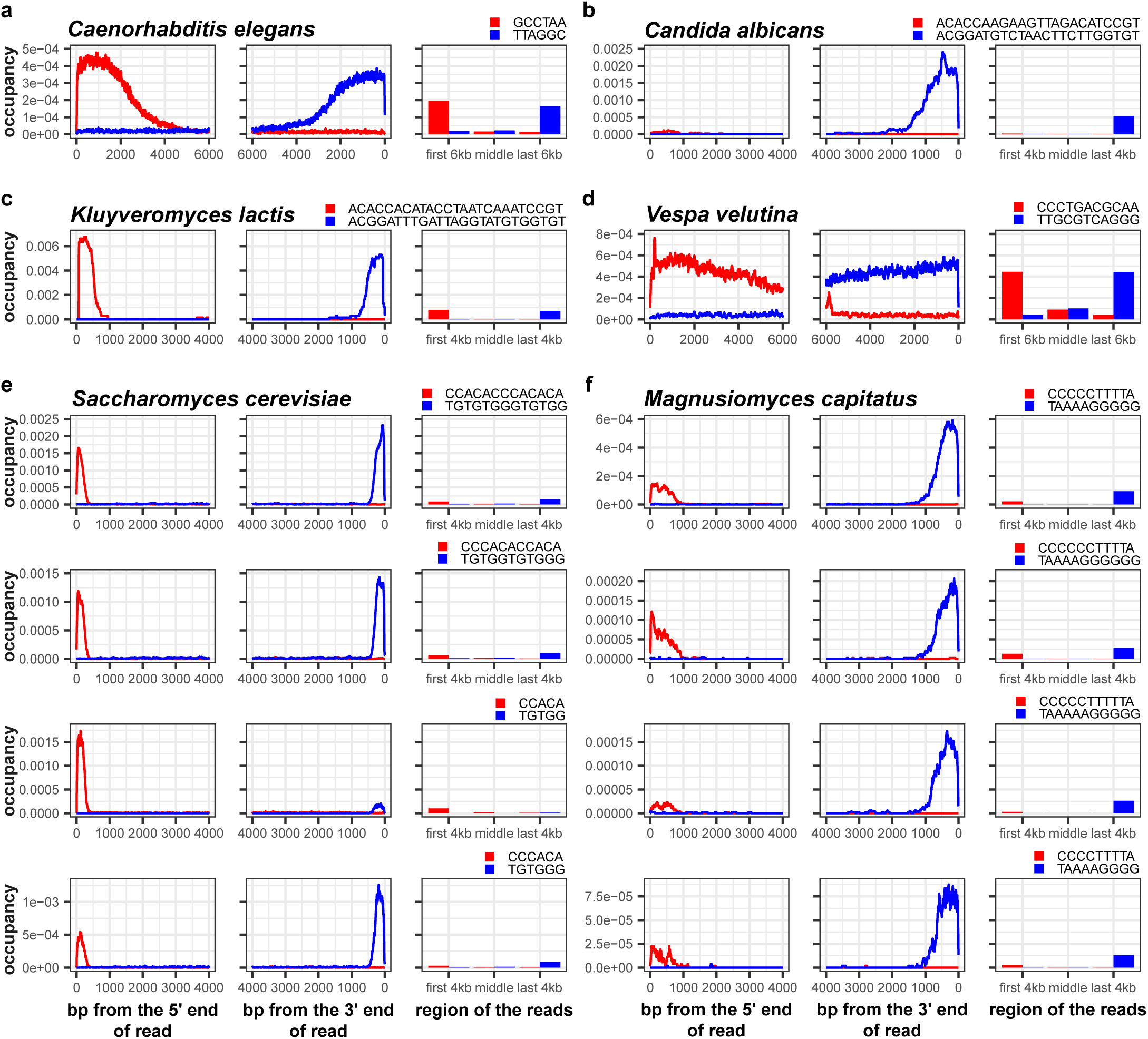
Detection of well-characterized telomere repeat motifs and variable telomeric repeat motifs. Occupancy patterns of TRMs (blue) and their reverse complements (red) for different species. **a,** TRM for the nematode *Caenorhabditis elegans*, TTAGGC. **b,** TRM for the yeast *Candida albicans,* ACGGATGTCTAACTTCTTGGTGT. **c,** TRM for the yeast *Kluyveromyces lactis*, ACGGATTTGATTAGGTATGTGGTGT. **d,** Top four most frequently observed TRMs for the brewing/baking yeast *Saccharomyces cerevisiae*. **e,** Top four TRMs for the yeast *Magnusiomyces capitatus*.

### Detection of telomeric motifs with variable repeat units

We next turned our attention to telomeric repeats motifs that are known to be variable. Several yeast species, including the widely studied *Saccharomyces cerevisiae* and the opportunistic pathogen *Magnusiomyces capitatus*, have telomeric repeat units that can vary in sequence and period due to the different possible ways to prime the extension of telomeric DNA using a telomerase RNA template^43^ or the stalling/stuttering of the telomerase enzyme during telomere lengthening^44^. Telomeric repeat motifs in *S. cerevisiae* and *M. capitatus* were previously reported to be (TG)_1–4_G_2-3_^45^ and T_1-2_A_3-5_G_3-8_^33^ respectively. To detect telomeric repeats from these two species, we ran TeloSearchLR in the occupancy mode to plot the occupancy of the 100 most frequently occurring 4-mer to 20-mer repeats in the terminal regions of sequencing reads (-k 4 -K 20 -m 1 -M 100). In both species, several top ranked terminal stranded occupancy patterns broadly matched the telomeric repeat motifs previously reported - (TG)_1–4_G_2-3_ for *S. cerevisiae* and T_1-2_A_3-6_G_4-6_ for *M. capitatus* (**Supplementary fig. 5-6**, red boxes). The occupancy plots of the top 4 repeat terminally stranded motifs are shown in **Fig. 2e-f**.

In *S. cerevisiae,* we further detected a stranded occupancy pattern for the sequence AGGGCTATTT and its reverse complement AAATAGCCCT, located consistently ∼400 bps from the ends of sequencing reads (**Supplementary fig. 5**, green box). Closer inspection revealed that these signals came from tandem repeats adjacent to the Y’ subtelomeric element in *S. cerevisiae*^46^. This demonstrated the potential for this algorithm to be used to detect subtelomeric tandem repeat patterns as well as telomeric tandem repeat patterns.

### Detection of coleopteran and hymenopteran telomeric repeat motifs missed by dot blot, Southern blot and short-read analysis

Confident that our algorithm could help us detect novel TRMs, we tested it on long sequencing reads from the insect orders Coleoptera (beetles) and Hymenoptera (bees, wasps, ants), where many species have divergent TRMs. In **Table 2** and **Figure 3**, we highlight 7 Coleopteran and Hymenopteran telomeric motifs missed by Southern blots, dot blots or short-read analysis^27–29^. Using occupancy plots generated by our algorithm, we detected stranded occupancy of the ancestral insect TTAGG telomeric motif in two species, *Diabrotica virgifera* and *Poecilus cupreus* (**Fig. 3a-b**, **Supplementary fig. 7-8**), but a previous report ruled out this ancestral motif^28^. We also successfully identified divergent TRMs in three genera (*Sitophilus oryzae* - TTTGG, *Geotrupes spiniger* - TTGGGG, and *Hyposoter dolosus* - TTTGGGTTTG) where hybridization-based assays and short-read analyses failed to detect TRMs in other members of the same genera (**Fig. 3c-e**, **Supplementary fig. 9-11**). The telomeric arrangement in *Geotrupes spiniger* is striking, as the telomeres appear to be only ∼50 bps long (**Fig. 3d**); however, other repeat motifs also show stranded and nearly terminal occupancy patterns (**Supplementary fig. 10**, green boxes), pointing to possible subtelomeric repeat sequences that may also cap the chromosome ends. Finally, for two species (*Diadromus collaris* and *Anthonomus grandis*), we could not detect any short motifs (4 to 50-mers) with clearly stranded occupancy patterns (**Supplementary fig. 12-13**).

**Figure 3:**
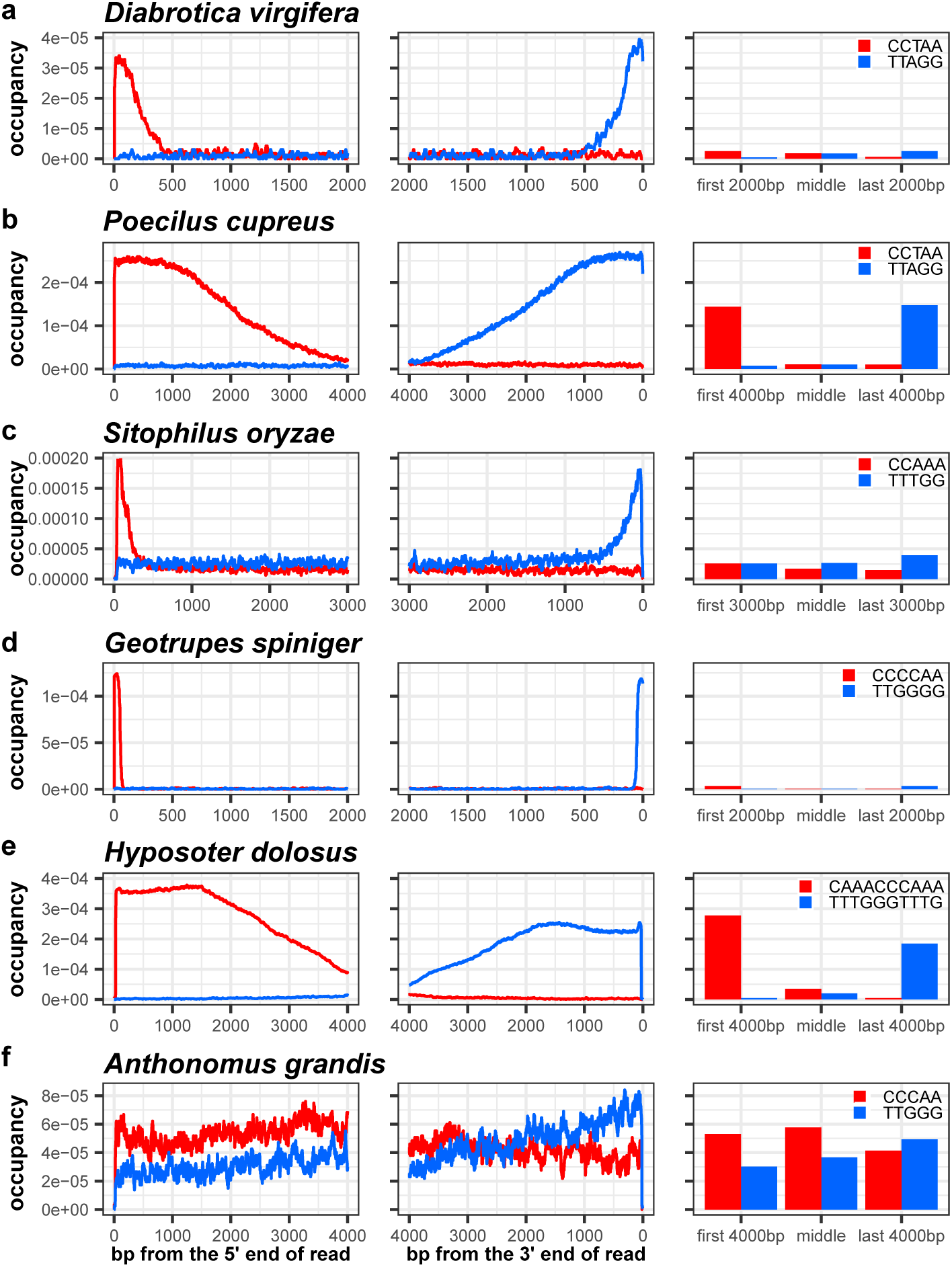
Detection of previously unidentified telomeric repeat motifs. Occupancy patterns of candidate TRMs (blue) and their reverse complements (red) for different species. **a,** Proposed TRM for the beetle *Diabrotica virgifera*, TTAGG. **b,** Proposed TRM for the ground beetle *Poecilus cupreus*, TTAGG. **c,** Proposed TRM for the rice weevil *Sitophilus oryzae*. **d,** Proposed TRM for the dung beetle *Geotrupes spiniger*, TTGGGG. **e,** Proposed TRM for the parasitoid wasp *Hyposoter dolosus*, TTTGGGTTTG. **f,** Candidate TRM for the boll weevil *Anthonomus grandis*, TTGGG. The occupancy patterns for this species do not show a heavy strand bias.

### Very long telomeres obscure the stranded occupancy of TRMs

Because our search strategy looks for motifs that are enriched at the 3’ or 5’ end of long sequencing reads, it would fail if the true telomeric repeat period were outside the range specified by the user, or if telomeres span the entire length of the reads. Thus, the *D. collaris* TRM may be longer than 50 bp. For *A. grandis*, we noticed that the occupancy patterns for TT<u>G</u>GG and its reverse complement CC<u>C</u>AA were slightly stranded (**Fig. 3f**), which suggested that the TRM evaded detection because the telomeres were much longer than the sequencing reads. When the sequencing reads are not long enough to capture entire telomeres, the counts of telomeric repeats would ‘spill over’ from the last *n* nucleotides to the middle and the first *n* nucleotides of the reads (**Fig. 4a**), obscuring the clear strandedness seen in **Fig. 1-3**. Similarly, the counts of the reverse complement patterns would ‘spill over’ to the last *n* nucleotides.

**Figure 4:**
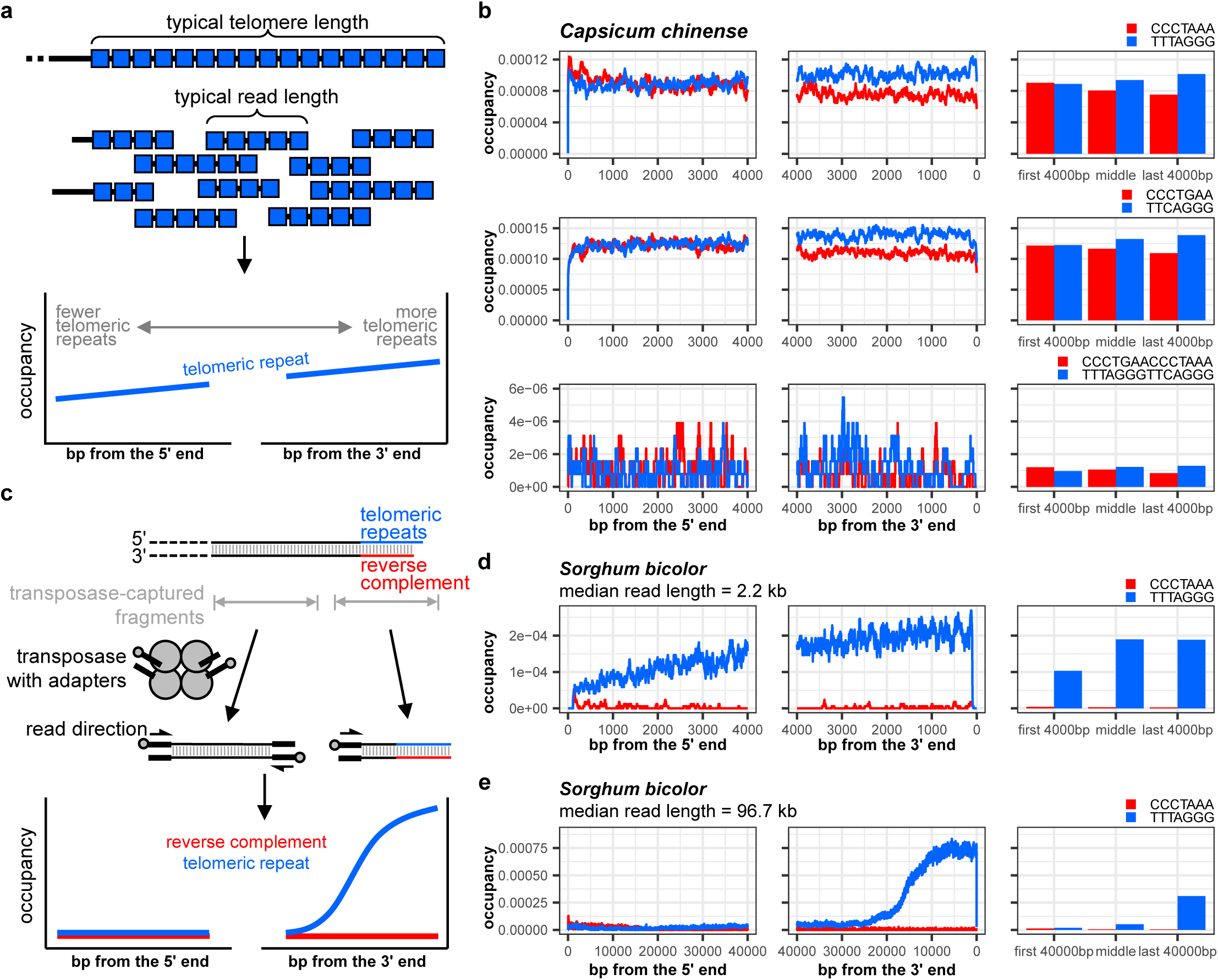
Known telomeric repeat motifs that evade detection. **a**, When the telomere is much longer than a typical sequencing read in a library, the telomeric repeat motif is no longer restricted to only one end of the read. Thus, the occupancy curve of the telomeric repeat does not show a stark stranded bias. The reverse complement is omitted for clarity. **b**, Occupancy plots of the *Capsicum chinense* telomeric repeat, TT[T/C]AGGG, with a variant third nucleotide. *C. chinense* telomeres have been shown to be maintained by telomerase and two paralogous *TERC* copies differing at the T/C variant position, and both TT<u>T</u>AGGG and TT<u>C</u>AGGG probes hybridize to the chromosome ends^46^. Thus, we looked for the occupancy patterns of TT<u>T</u>AGGG, TT<u>C</u>AGGG, and TT<u>T</u>AGGGTT<u>C</u>AGGG repeats. Their occupancy patterns do not appear to be markedly stranded. **c**, One method of preparing “ultra-long” Nanopore sequencing library involves using a specialized transposase optimized for the attachment of sequencing adapters 50-100kb apart. Near the chromosome end, it may not be possible to attach both adapters, leading to telomeric reads favoring the orientation with the telomeric repeats at the 3’ end. **d**, Occupancy plots of the *Sorghum bicolor* telomeric repeat, TTTAGGG, for a library with a median read length of 2.2 kb. **e**, Occupancy plots of the *S. bicolor* telomeric repeat, TTTAGGG, for an “ultra-long” sequencing library with a median read length of 96.7 kb.

Supporting this idea, known TRMs sometimes do not have a clear terminal stranded occupancy pattern on long sequencing reads (**Fig. 4a**). For example, the telomeric repeat motif of the Habanero pepper *Capsicum chinense*, previously found to be TT[T/C]AGGG^47^, does not have an obviously stranded occupancy with the plotting window *n* = 4 kb, in a long-read library with a median read length of 13.5 kb (run accession SRR23734611) (**Table 2**, **Fig. 4b** and **Supplementary fig. 14**). To see if the stranded occupancy pattern could be revealed by extending the plotting windows, we adjusted the *n* value from 4,000 to 11,000 (**Supplementary fig. 15a, c-j**). However, this also had the effect of decreasing the number of reads being considered (**Supplementary fig. 15b**), as fewer reads were 22,000 bps or longer (2 × 11,000 bps) in comparison with reads that were 8,000 bps or longer (2 × 4,000 bps) (**Supplementary fig. 15b**). We could only detect a terminal stranded occupancy with *n* = 9000 or above (**Supplementary fig. 15h-j**), but the median telomere length could no longer be estimated since there were too few telomeric reads 18,000 bps or longer. This result indicates that *C. chinense* telomeres are often 9000 bp or longer. Our search strategy thus depends on genomic sequencing reads being much longer than typical telomeres in a biological sample.

### Detection of TRMs from long telomeres using ultra-long sequencing reads

Genome sequencing library preparation methods exist to generate “ultra-long” sequencing reads with lengths of up to 50 to 100 kb using specialized transposases to attach sequencing adapters and their pre-loaded sequencing helicases (**Fig. 4c**). For naturally long (>10kb) telomeres, these ultra-long reads might reveal the terminal stranded occupancy patterns of the TRM. We found that the sorghum (*Sorghum bicolor*) motif, TTTAGGG^48^, did not have a typical terminal stranded occupancy from a library with a median read length of 2.2 kb (run accession ERR4312692) (**Table 2**, **Fig. 4d**, and **Supplementary fig. 16**). However, in an ultra-long genomic library (run accession SRR26636576) with a median read length of 96.7 kb, we found the TTTAGGG telomeric repeat motif at the 3’ ends of the reads, but comparatively few CCCTAAA reverse complement repeats at the 5’ ends (**Fig. 4e**, **Supplementary fig. 17**), with an estimated telomere median length of 15 kb.

The bias for telomeric repeats on the 3’ end may be due to the way these libraries were generated. Near the chromosome end, the transposase may attach a sequencing adapter only at the proximal end and not the distal end, given that the transposase has been engineered for attaching adapters a specific distance apart (**Fig. 4c**). Thus, these ultra-long telomeric reads would heavily favor having telomeric repeats at the 3’ ends, rather than with the telomeric reverse complement at the 5’ ends of the reads. Thus, the occupancy plots would not be mirrored, but instead show a more prominent occupancy curve for the 3’ telomeric motif than for its 5’ reverse complement (**Fig. 4c** and **e**), unlike most of the instances we have examined so far (**Fig. 1-3**).

### Detection of telomeric motifs with unusually long repeat periods maintained by telomerase-independent mechanisms

So far, we have focused on organisms that lengthen and maintain telomeres using telomerase – the predominant eukaryotic strategy to deal with the end replication problem; however, the telomerase gene has been lost in several eukaryotic lineages, with the most studied example being the Dipteran order (flies), where specialized transposons maintain telomere length^49^. These telomerase-independent TRMs can be much longer than TRMs associated with the use of telomerase. Extending our algorithm to search for telomeres maintained by telomerase-independent mechanisms, we allowed the ranking step of TeloSearchLR to examine more than 1000 bp. This in turn allowed the algorithm to examine tandem repeats longer than 500 bp (as the longest *tandem* repeat motif to fit inside 1000 bp is 500 bp).

We chose to test our algorithm on the sequencing reads from the parasitic nematode *Strongyloides stercoralis*, for which a previous attempt to identify telomere repeat motifs using a short-read search strategy had been unsuccessful^36^. Due to the simple karyotype of 2*n* = 6, we reasoned *S. stercoralis* was the ideal organism to test, as there would only be 6 distinct chromosome ends to resolve in a homozygote. In addition, we were aided by a very contiguous reference genome assembly available for this species (GenBank accession GCA_029582065.1), with two autosomal scaffolds and several X chromosome scaffolds. It was unclear if the scaffold ends were fully resolved to reveal the true telomeres, so we aimed to identify all six telomeric ends of this species.

Our search strategy did not turn up any short repeat motifs (4-20 bps) that had a terminal stranded occupancy (-k 4 -K 20 -t 1000, **Supplementary fig. 18**), so we examined tandem repeat motifs up to 1000 bps long, ranked by occupancies in the terminal 2000 bps on reads longer than 4000 bps (-k 21 -K 1000 -t 2000). Out of the top 100 repeat motifs (-m 1 -M 100), three repeat motifs and their reverse complements had terminal stranded occupancy, albeit with a noticeable spillover effect like *A. grandis* or *C. chinense*, indicating that these telomeres are long. These three motifs are repeat pattern #6 (a 111-mer), #7 (a 106-mer), and #9 (a 105-mer) (**Fig. 5a-c**, **Supplementary fig. 19**). The three motifs share significant sequence similarity and are likely to have descended from a common sequence (**Fig. 5d**). We used the TideHunter output to extract reads with patterns #6, #7 and #9 and found that they mapped to the assembled scaffold ends (**Fig. 5e**, **Supplementary fig. 22a, c**, and see **Supplementary Notes**). Thus, we propose that these scaffold ends are the true telomeres of *S. stercoralis*, and we propose that the two X chromosome telomeres are located on X chromosome scaffolds X2 and X3.

**Figure 5:**
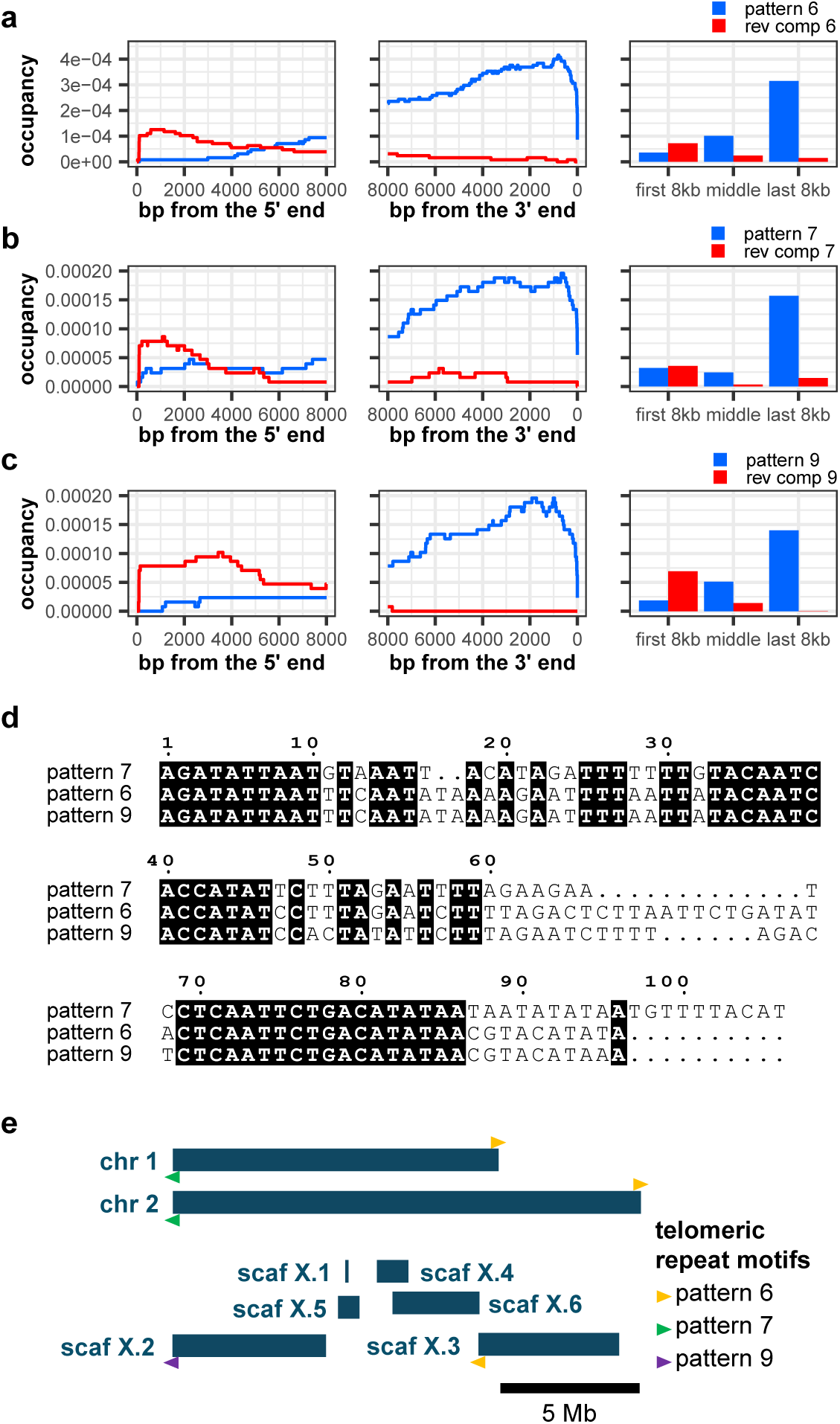
Telomeric repeat motifs of *Strongyloides stercoralis*. Occupancy patterns of top three *S. stercoralis* repeat motifs with stranded bias (blue) and their reverse complements (red). **a,** Repeat motif #6, a 111-mer. **b,** Repeat motif #7, a 106-mer. **c,** Repeat motif #9, a 105-mer. **d,** Multiple sequence alignment of motifs 6, 7 and 9. **e,** The locations of telomeres with motif 6 (yellow), 7 (green) and 9 (purple) in the genome assembly GCA_029582065.1. Arrowheads above the chromosome or scaffold schematic indicate the telomeric motifs in their conventional orientation (pointing toward the end of the chromosome), while those below indicate motifs represented by their reverse complement in the assembly. Scaffolds X.2 and X.3 are likely terminal scaffolds of the X chromosome.

### Validation of telomere length estimates

We compared several of our telomere length estimates with published estimates to see if ours were in line with estimates based on Southern blots and various short-read repeat analyses (**Table 3**). This comparison is not perfect because the biological isolates used for long-read sequencing libraries are not identical to those used in separate studies. Nevertheless we found that, except for *C. elegans*, our telomere length estimates were broadly in line with previous estimates.

**Table 3:**
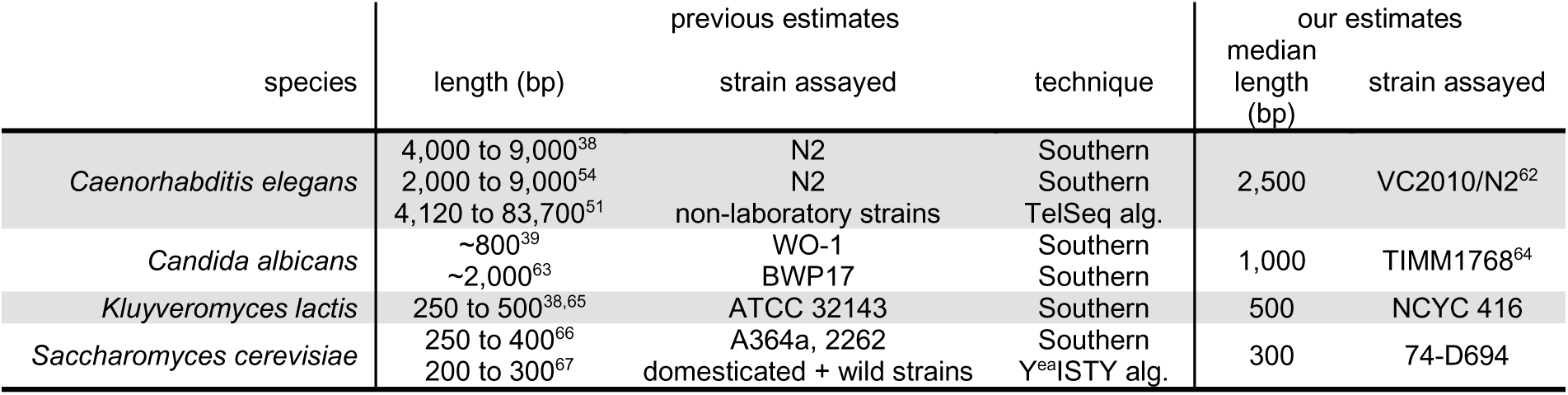
Comparison of telomere length estimates

## DISCUSSION

We developed a telomere motif search strategy guided by TeloSearchLR, an intuitive algorithm whose graphical output allows the visual identification of telomeric repeat motifs at the ends of long sequencing reads (**Table 2**). We demonstrated that our search strategy was compatible with sequencing reads from both Pacific Biosciences (PacBio) or Oxford Nanopore Technologies (ONT) platforms (**Table 2**), and that our strategy complemented existing telomere motif searches by successfully identifying novel TRMs where they were missed (**Table 2**). Testing on a variety of different sequencing libraries, we found that our strategy succeeded when sequencing reads were much longer than typical telomeres in a biological sample. Moreover, we demonstrated that the median telomere length could be inferred directly from the occupancy plots generated by our algorithm.

### The use of stranded repeat motif position allows the detection of TRMs from naturally short telomeres

Short-read TRM search strategies focus on the frequency of repeat motifs. In addition to frequency, TeloSearchLR also considers the position of the repeat motif occurrences, which is possible because long sequencing reads preserve natural DNA ends. Thus, an advantage to our approach over short-read search strategies is our ability to identify TRMs from naturally short telomeres, such as those from *D. virgifera* (median ≈ 200 bp). We suspect that the *D. virgifera* TRM eluded detection by short-read searches because the TTAGG motif and its reverse complement were not heavily enriched in the sequencing library^28^. In our analysis, TTAGG and its reverse complement represent only the 54th most frequent motif in the terminal 1000 bp of the *D. virgifera* sequencing reads (**Supplementary fig. 7**), much lower ranked than TRMs in other species. Nevertheless, using TeloSearchLR we were able to conclude that TTAGG was the *D. virgifera* TRM by its terminal stranded occupancy on the sequencing reads. Thus, the consideration of repeat motif position on sequencing reads by TeloSearchLR provides a worthy addition to existing TRM search tools.

### TRMs from very long telomeres can be detected on even longer sequencing reads

When telomeres are of comparable or greater length than available sequencing reads, telomeric repeats are not enriched at opposite ends of the sequencing reads (**Fig. 4a**). We demonstrated that this limitation to our search strategy could be overcome partly by using sequencing libraries specifically designed to yield very long and intact DNA (**Fig. 4e**). In our analysis of the *S. bicolor* telomeres, a library with a median read length of 95 kb revealed the telomeres with a median length of 15 kb. Ultra-long reads also benefit genome assembly efforts as well – so that genome assemblers can fully resolve telomeric and other repetitive regions. Thus, we recommend the use of the longest possible sequencing read lengths for any novel sequencing project.

### TeloSearchLR can be used to anchor terminal scaffolds in new genome assemblies

Our strategy for identifying telomeric repeat motifs also complements new genome assembly efforts by anchoring terminal scaffolds or contigs to their respective chromosome ends. The analysis of *Strongyloides stercoralis* sequencing reads revealed specific 105-bp, 106-bp and 111-bp tandem repeats at the ends of sequencing reads, identical to the repeats at the ends of the current genome assembly of *S. stercoralis* (GenBank accession GCA_029582065.1). We therefore propose that TeloSearchLR may be used as a quality checking step in the assembly of new genomes: when chromosome ends are fully resolved they should include telomeres revealed by TeloSearchLR.

### Telomere lengths estimated from repeat occupancy plots may be shorter than other estimates

We expect our method to produce slightly shorter telomere length estimates compared to Southern blots. This is because our telomeric repeat occupancy tallies use ‘noisy’ reads that are not completely error-free. While the TideHunter tandem repeat detection algorithm tolerates some sequencing errors^37^, manual inspection of the TideHunter output revealed that telomeric repeat units with too many mismatches (likely from sequencing errors) were not counted (**Supplementary fig. 20a-b**). This undercounting would lead to a shorter telomere length estimate (**Supplementary fig. 20c**), as telomeres would appear to have fewer telomeric repeat units.

DNA library preparation methods also contribute to the underestimation of telomere length. Our source data come from typical genomic sequencing libraries that use end-blunting enzymes (and an A-tailing enzyme) to facilitate the attachment of double-stranded sequencing adapters (**Supplementary fig. 21a**). These procedures are expected to eliminate 3’ single-stranded overhangs at telomeres. As a result, our algorithm only surveys the double-stranded portion of the telomere, whereas Southern blots can reveal the single-stranded portion when it is being probed.

### A biased orientation places telomeric repeats at the 3’ end of sequencing reads

We initially expected the occupancy pattern of the TRM to mirror that of its reverse complement on the opposite ends of sequencing reads in all of the libraries we analyzed (**Fig. 1a**). However, motif occupancy patterns in some libraries are not mirrored, and most of these show a stronger preference for TRMs at the 3’ end of reads. We infer that technical differences in library preparation protocols, combined with specific characteristics of chromosome ends in some species, can give rise to biased attachment of sequencing adaptors to either end of reads derived from telomeric sequences. For example, for the *C. albicans* (**Fig. 2b**) and *M. capitatus* (**Fig. 2d**) libraries, we hypothesize that non-B-DNA telomeric structures such as T-loops^50^ and guanine quadruplexes^51,52^ are more resistant to DNA blunting and adapter ligation than standard B-form double-stranded DNA (**Supplementary fig. 21b-c**). In the case of *S. bicolor* (**Fig. 4c**), we attribute the asymmetry in motif end enrichment to the use of a transposase during library preparation. All of these examples would lead to the preferential attachment of sequencing adapters at the non-telomeric end of terminal DNA fragments, leading to a loss of reads enriched for the reverse complement of TRMs at their 5’ end and thus preferential recovery of reads with TRMs at the 3’ end.

### Extension of the algorithm to detect repetitive subtelomeres

Even though our method was specifically designed to detect telomeric repeat motifs, our analysis of the *S. cerevisiae* reads revealed a tandem repeat adjacent to the Y’ subtelomeric element. Similarly, we discovered several stranded occupancy patterns that were not terminal in *G. spiniger* but which could represent subtelomeric repetitive sequences (**Supplementary fig. 10**, green boxes). This means that subtelomeric sequences, if they contain tandem repeats, can also be identified by our method.

In conclusion, we demonstrated that a novel telomere search strategy using TeloSearchLR could successfully identify novel telomeric repeats while providing an estimate of telomere length. When used in conjunction with genome assembly, TeloSearchLR provides a means to anchor telomeric genome scaffolds and to check the assembly ends. Moreover, our algorithm can be extended to identify subtelomeric repetitive elements. TeloSearchLR thus provides a valuable addition to existing tools for identifying telomere-associated repeat motifs.

## WEB RESOURCES AND DATA AVAILABILITY

The TeloSearchLR.py source code is available via the TeloSearchLR repository on GitHub (https://github.com/gchchung/TeloSearchLR).

## ACKNOWLEDGEMENTS

The authors wish to thank Dr. Karin Kiontke for her suggestions on the manuscript. The authors would also like to thank Dr. Andreas Hochwagen and Adhithi Raghavan for their insight on *S. cerevisiae* subtelomeric repeats.

## SUPPLEMENTARY NOTES

### Mapping *Strongyloides stercoralis* telomeric reads to the assembly

In the *Strongyloides stercoralis* sequencing libraries (SRA run accessions SRR25177361 and SRR25177362), the three putative telomeric motifs found by TeloSearchLR are present at five of the six assembled chromosome ends in *S. stercoralis* (GenBank accession: GCA_029582065.1). The three motifs are named pattern 6, pattern 7 and pattern 9. The only chromosome end without one of these patterns was the right end of chr 1. This meant that the right end of chr 1 could be capped by a different (and potentially non-repetitive) sequence, or that this end was assembled incompletely or incorrectly.

To test if the right end of chr 1 was assembled correctly, we isolated reads with pattern 6, 7, and 9 repeats and mapped them to the assembly using minimap2^53^. Because TeloSearchLR results provided compelling evidence these were telomeric reads, any of these mapping to or near the right end of chr 1 could help us identify the true chromosome end. Reads with pattern 7 repeats mapped to the left ends of chr 1 and chr 2 (**Supplementary fig. 22a**). Reads with pattern 9 repeats mapped to the left end of scaffold X.2 (**Supplementary fig. 22a**). Reads with pattern 6 repeats mapped to the left end of scaffold X.3, the right end of chromosome 2, and to an interstitial region on chr 1, where alignments ended abruptly at position 11,637,827 roughly 35 kb from the assembled right end (**Supplementary fig. 22a**, arrowhead). For sequencing reads that map to this last location, instead of having parts of the ∼35 kb sequence found in the assembly, these reads had a ∼1490 bp of subtelomeric sequence before terminating in tandem repeats of pattern 6 (**Supplementary fig. 22b**). Thus, the ∼35-kb sequence (11,637,828 - 11,673,123) at the right end of chr 1 was misassembled, and in its place should be the missing 1490-bp subtelomere and tandem repeats of pattern 6 as the telomere (**Supplementary fig. 22b**). We re-aligned the sequencing reads with pattern 6 repeats to this corrected configuration and found that they supported this chromosome end structure (**Supplementary fig. 22c**).

**Supplementary figure 1: Occupancy plots of the 100 most frequently appearing 4- to 20-mer terminal repeat motifs in a *Caenorhabditis elegans* genomic sequencing library.**

The occupancy pattern for the *C. elegans* telomeric repeat motif, TTAGGC, is highlighted by the red box, while patterns for several telomere-repeat-like motifs are highlighted in pink.

**Supplementary figure 2: Occupancy plots of the 100 most frequently appearing 4- to 30-mer terminal repeat motifs in a *Candida albicans* genomic sequencing library.**

The occupancy pattern for the *C. albicans* telomeric repeat motif, ACGGATGTCTAACTTCTTGGTGT, is highlighted by the red box, while patterns for several telomere-repeat-like motifs are highlighted in pink.

**Supplementary figure 3: Occupancy plots of the 100 most frequently appearing 4- to 30-mer terminal repeat motifs in a *Kluyveromyces lactis* genomic sequencing library.**

The occupancy pattern for the *C. albicans* telomeric repeat motif, ACGGATTTGATTAGGTATGTGGTGT (as the cyclical equivalent ATTTGATTAGGTATGTGGTGTACGG), is highlighted by the red box.

**Supplementary figure 4: Occupancy plots of the 100 most frequently appearing 4- to 20-mer terminal repeat motifs in a *Vespa velutina* genomic sequencing library.**

The occupancy pattern for the *V. velutina* telomeric repeat motif, TCAGGGTTGCG (as the cyclical equivalent GCGTCAGGGTT), is highlighted by the red box. The pattern for a telomere-like motif with 1 nucleotide difference is highlighted in pink.

**Supplementary figure 5: Occupancy plots of the 100 most frequently appearing 4- to 20-mer terminal repeat motifs in a *Saccharomyces cerevisiae* genomic sequencing library.**

The occupancy patterns for the *S. cerevisiae* telomeric repeat motifs are highlighted by the red boxes. The stranded occupancy pattern of a tandem repeat (AGGGCTATTT) from the Y’ subtelomeric element is highlighted in green.

**Supplementary figure 6: Occupancy plots of the 100 most frequently appearing 4- to 20-mer terminal repeat motifs in a *Magnusiomyces capitatus* genomic sequencing library.**

Occupancy patterns for the *M. capitatus* telomeric repeat motifs are highlighted by the red boxes.

**Supplementary figure 7: Occupancy plots of the 100 most frequently appearing 4- to 20-mer terminal repeat motifs in a *Diabrotica virgifera* genomic sequencing library.**

The occupancy pattern for the *D. virgifera* telomeric repeat motif candidate, TTAGG, is highlighted by the red box.

**Supplementary figure 8: Occupancy plots of the 100 most frequently appearing 4- to 20-mer terminal repeat motifs in a *Poecilus cupreus* genomic sequencing library.**

The occupancy pattern for the *P. cupreus* telomeric repeat motif candidate, TTAGG, is highlighted by the red box. Patterns for possible subtelomeric repeat motifs are highlighted in green.

**Supplementary figure 9: Occupancy plots of the 100 most frequently appearing 4- to 20-mer terminal repeat motifs in a *Sitophilus oryzae* genomic sequencing library.**

The occupancy pattern for the *S. oryzae* telomeric repeat motif candidate, TTTGG, is highlighted by the red box.

**Supplementary figure 10: Occupancy plots of the 100 most frequently appearing 4- to 20-mer terminal repeat motifs in a *Geotrupes spiniger* genomic sequencing library.**

Plotting window is 4 kb – unlike in Fig. 3d – to reveal occupancy patterns of possible subtelomeric repeat motifs. The occupancy pattern for the *G. spiniger* telomeric repeat motif candidate, TTGGGG, is highlighted by the red box. Occupancy patterns for possible subtelomeric repeat motifs – AACGGCGGGTGCGCGGGTA and TACCCA – are highlighted in green.

**Supplementary figure 11: Occupancy plots of the 100 most frequently appearing 4- to 20-mer terminal repeat motifs in a *Hyposoter dolosus* genomic sequencing library.**

The occupancy pattern for the *H. dolosus* telomeric repeat motif candidate, TTTGTTTGGG, is highlighted by the red box.

**Supplementary figure 12: Occupancy plots of the 40 most frequently appearing 4- to 50-mer terminal repeat motifs in a *Diadromus collaris* genomic sequencing library, grouped by repeat period.**

**Supplementary figure 13: Occupancy plots of the 40 most frequently appearing 4- to 50-mer terminal repeat motifs in an *Anthonomus grandis* genomic sequencing library, grouped by repeat period.**

**Supplementary figure 14: Occupancy plots of the 100 most frequently appearing 4- to 20-mer terminal repeat motifs in a *Capsicum chinense* genomic sequencing library.**

The occupancy patterns for the two known *C. chinense* telomeric repeat motifs, TT**C**AGGG and TT**T**AGGG, are highlighted by grey boxes.

**Supplementary figure 15: Effects of increasing the plotting window length *n* on occupancy plots of repeat motifs in a *Capsicum chinense* genomic sequencing library.**

**a**, Graphical representation of changing the graphing window value *n*. The sequencing reads must be longer than or equal to 2*n* nucleotides long for the repeat occupancies counted. **b**, Distribution of sequencing read lengths for the sequencing library SRR23734611, with the y-axis in log scale to reveal the few very long reads. As the value of *n* increases, fewer sequencing reads will be 2*n* nucleotides or longer. **c**, The occupancy patterns of the two *C. chinense* telomeric repeat motifs, TT**C**AGGG and TT**T**AGGG, with a plotting window of *n* = 4000 bps. **d-j**, The occupancy patterns with *n* = 5000, 6000, 7000… 10000, 11000 bps. A terminal stranded occupancy pattern is apparent starting from *n* = 9000 (**h**).

**Supplementary figure 16: Occupancy plots of the 100 most frequently appearing 4- to 20-mer terminal repeat motifs in a *Sorghum bicolor* genomic sequencing library.**

The occupancy pattern for the *S. bicolor* telomeric repeat motif candidate, TTTAGGG, is highlighted by the grey box.

**Supplementary figure 17: Occupancy plots of the 100 most frequently appearing 4- to 20-mer terminal repeat motifs in a *Sorghum bicolor* ultra-long-read genomic sequencing library.**

The occupancy pattern for the *S. bicolor* telomeric repeat motif candidate, TTTAGGG, is highlighted by the red box. A strongly stranded occupancy pattern is only detected in the ultra-long-read data (compare vs. grey box in Supplementary fig. 16).

**Supplementary figure 18: Occupancy plots of the 100 most frequently appearing 4- to 20-mer terminal repeat motifs in a *Strongyloides stercoralis* genomic sequencing library.**

**Supplementary figure 19: Occupancy plots of the 100 most frequently appearing 21- to 1000-mer terminal repeat motifs in a *S. stercoralis* genomic sequencing library.**

**Supplementary figure 20: Sequencing errors in the telomeric motif can lead to under-counting of repeat numbers and a shorter telomere length estimate.**

**a,** An idealized depiction of telomeric reads with the telomeric reverse complement motifs at the 5’ ends of the reads. Red boxes indicate the position and the span of a telomeric repeat unit. **b,** In reality, not all of the repeat units are counted due to sequencing errors, indicated by the green ‘x’. **c,** As a result, median estimates of the telomeric length may be shorter (green) than the true median (red) due to noisy reads and under-counting of telomeric repeat units.

**Supplementary figure 21: Typical long-read sequencing library construction and telomeric reads with sequencing strand bias.**

**a,** Sequencing library construction involves enzymatic blunting and A-tailing to the DNA to be sequenced. In the blunting step, the usual 3’ telomeric overhang is expected to be deleted by the blunting enzyme.

Thus, the single-stranded telomere sequence is lost in the sequencing step, leading to a shorter telomere length estimate by our method. **b,** The G-rich telomeric strand (blue) is prone to form guanine quadruplex structures under certain conditions. This may render the telomere end resistant to end repair and sequencing adapter attachment, leading to a sequencing strand orientation bias. **c,** The 3’ telomeric overhang can form T-loops. This may render the telomere end resistant to end repair and sequencing adapter attachment, leading to a sequencing strand orientation bias.

**Supplementary figure 22: *Strongyloides stercoralis* sequencing reads with pattern 6, 7 and 9 repeats map to assembly ends.**

**a,** *S. stercoralis* genomic sequencing reads (SRA run accessions: SRR25177361 and SRR25177362) with pattern 6, 7 and 9 tandem repeats map to five assembly ends (GenBank accession GCA_029582065.1): reads with pattern 6 repeats map to chr 2 right and scaffold X.3 left; reads with pattern 7 repeats map to chr 1 left and chr 2 left; and reads with pattern 9 repeats map to scaffold X.2 left. Pattern 6 reads also map to a region near the right end of chr 1, but the alignments abruptly end at position 11,637,827, which suggests a misassembly event. The bottommost track indicates the location and the type of tandem repeats at the ends of the assembly. **b,** Sequencing reads with pattern 6 repeats that map to chr 1: 11,637,827 do not agree with the assembled sequence. Instead of having parts of the ∼35-kb sequence from 11,647,828 bp to the end of the assembly, the reads have a ∼1490-bp subtelomeric sequence followed by tandem repeats of pattern 6. **c,** Sequencing reads with pattern 6 repeats support the corrected chr 1 end with the ∼1490-bp sequence and pattern 6 tandem repeats.

