## Supplementary figures and images for "TeloSearchLR: an algorithm to detect novel telomere repeat motifs using long sequencing reads"

### Supp. fig. 1

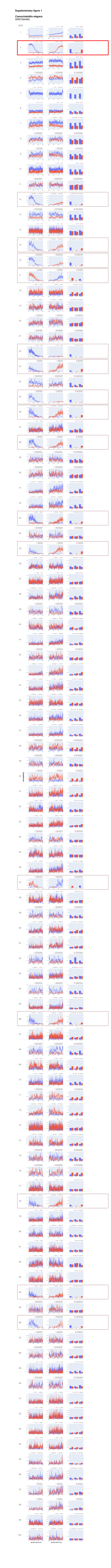

### Supp. fig. 2

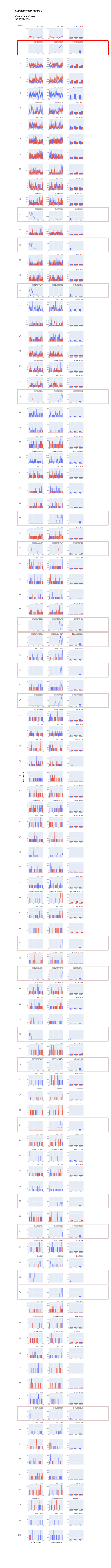

### Supp. fig. 3

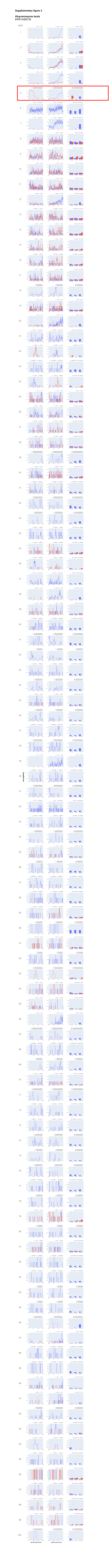

### Supp. fig. 4

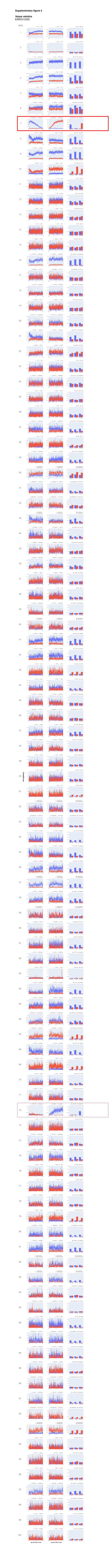

### Supp. fig. 5

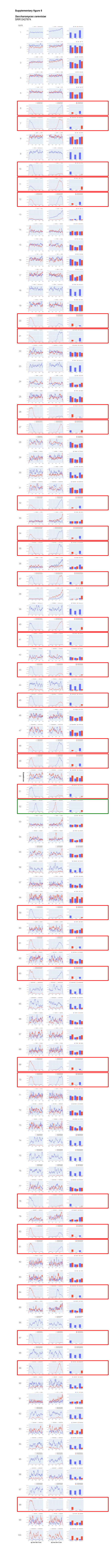

### Supp. fig. 6

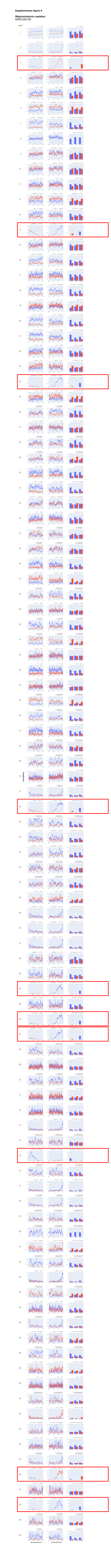

### Supp. fig. 7

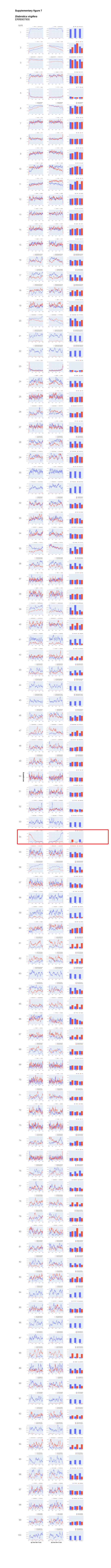

### Supp. fig. 8

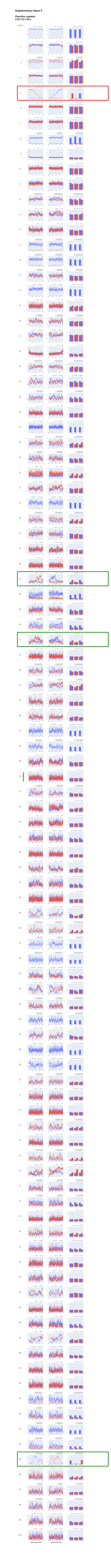

### Supp. fig. 9

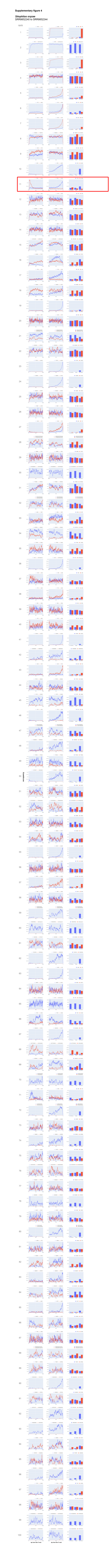

### Supp. fig. 10

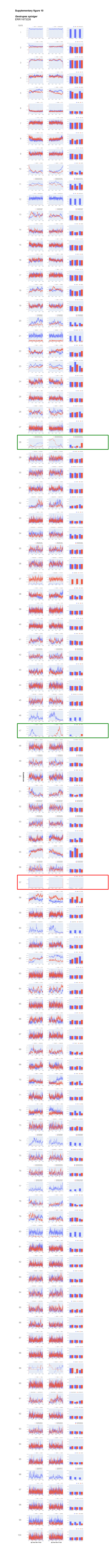

### Supp. fig. 11

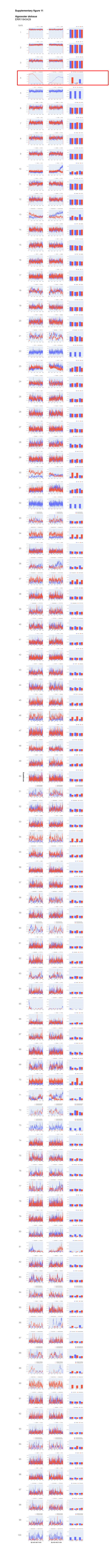

### Supp. fig. 14

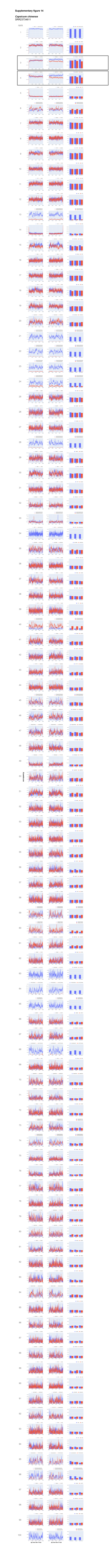

### Supp. fig. 15

#### Supplementary figure 15

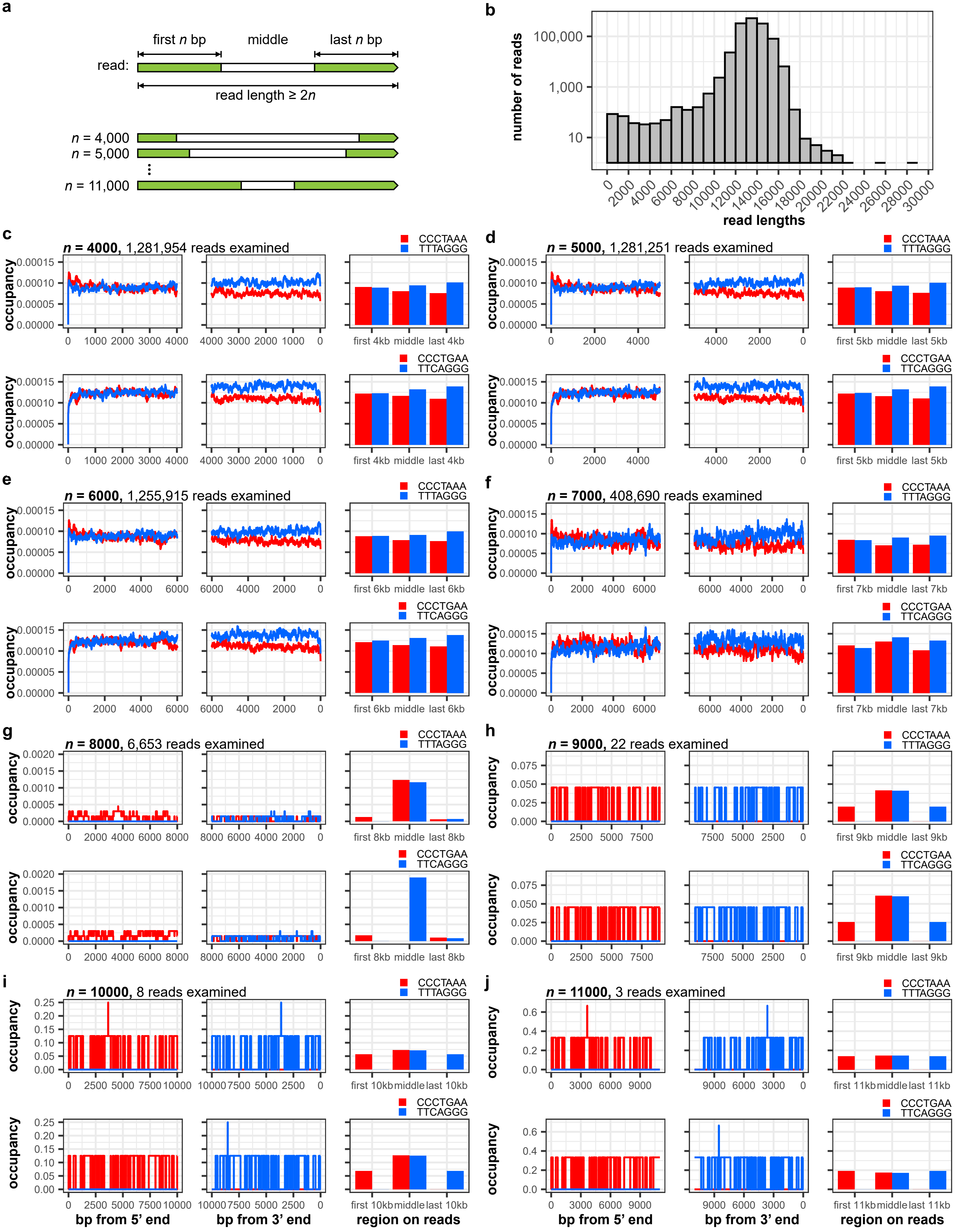

### Supp. fig. 16

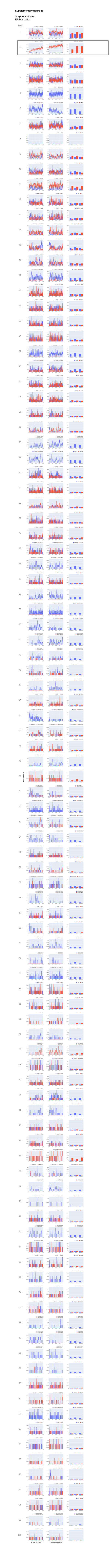

### Supp. fig. 17

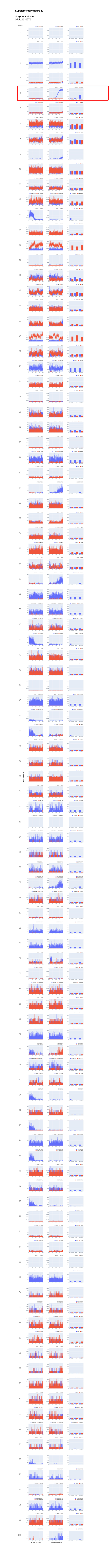

### Supp. fig. 18

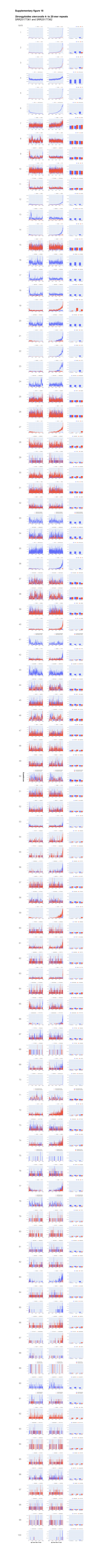

### Supp. fig. 19

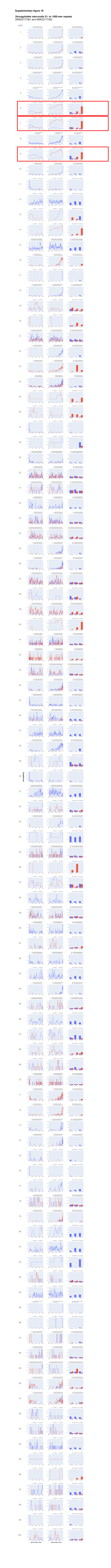

### Supp. fig. 20

#### Supplementary figure 20

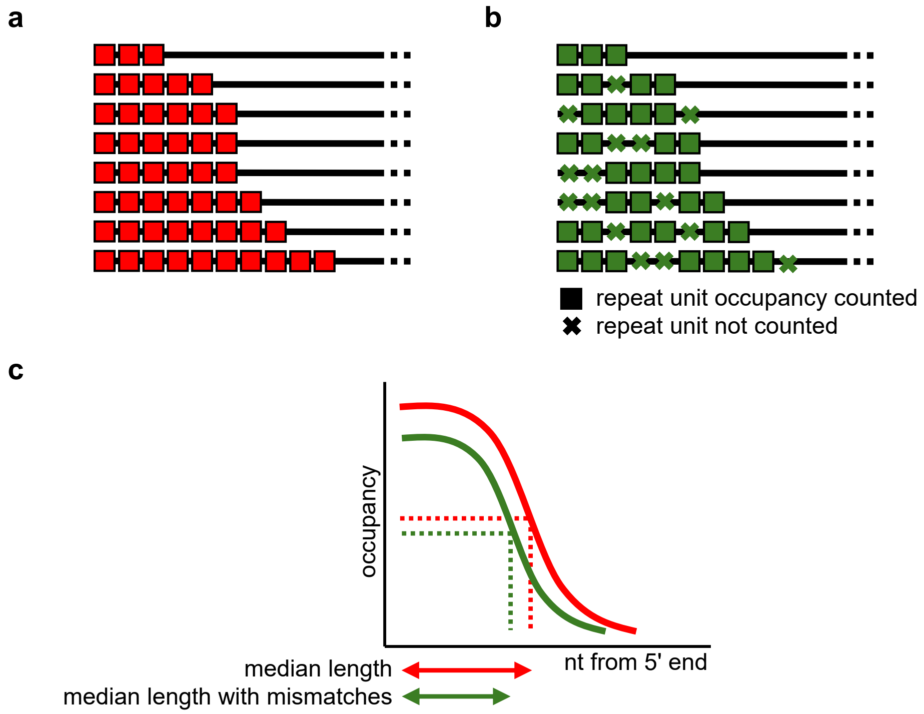

### Supp. fig. 21

**Supplementary figure 21**

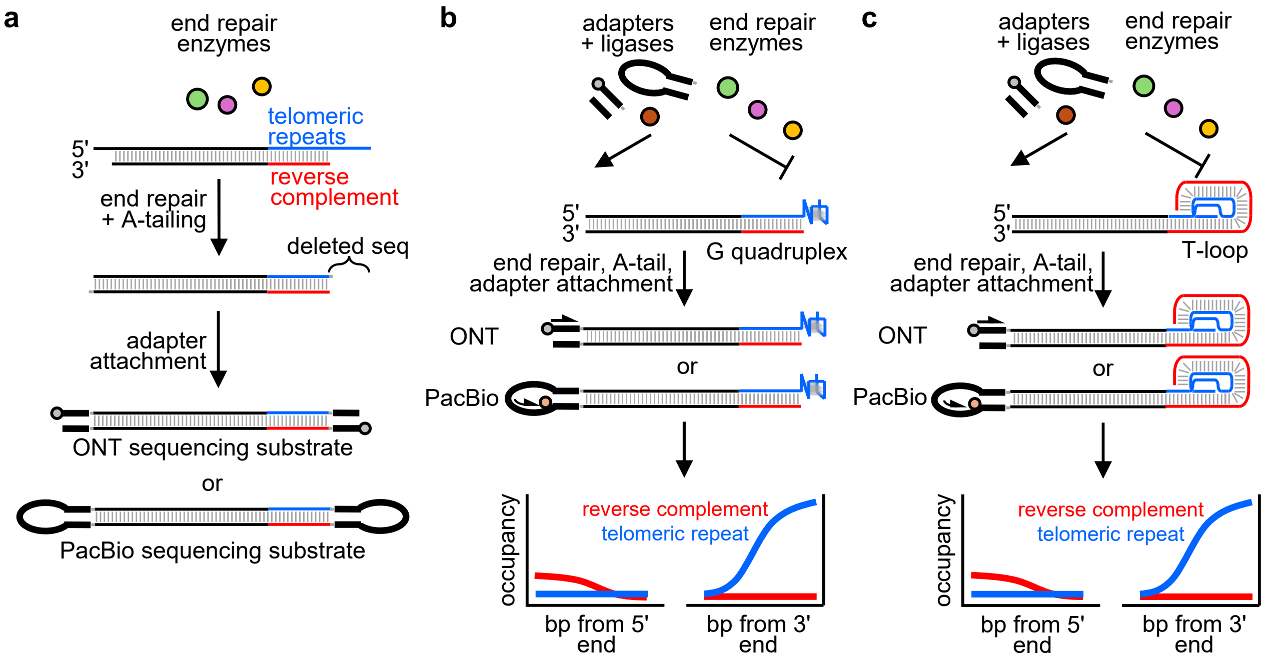
