## Supplementary material for "TeloSearchLR: an algorithm to detect novel telomere repeat motifs using long sequencing reads": Supp. fig. 22

**Supplementary fig. 22**

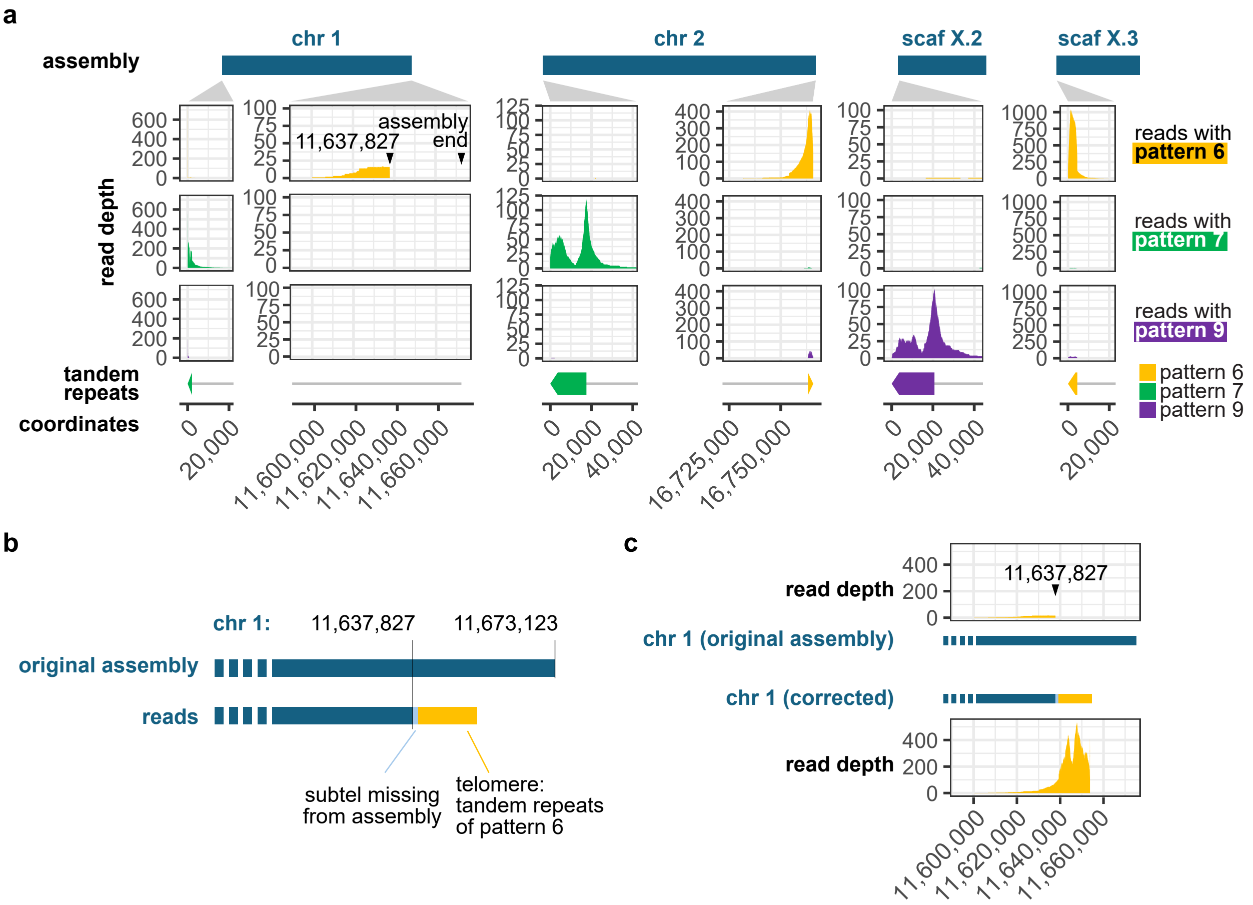


**Supplementary figure 22: *Strongyloides stercoralis* sequencing reads with pattern 6, 7 and 9 repeats map to assembly ends.**

**a,** *S. stercoralis* genomic sequencing reads (SRA run accessions: SRR25177361 and SRR25177362) with pattern 6, 7 and 9 tandem repeats map to five assembly ends (Genbank accession GCA_029582065.1): reads with pattern 6 repeats map to chr 2 right and scaffold X.3 left; reads with pattern 7 repeats map to chr 1 left and chr 2 left; and reads with pattern 9 repeats map to scaffold X.2 left. Pattern 6 reads also map to a region near the right end of chr 1, but the alignments abruptly end at position 11,637,827, which suggests a misassembly event. **b,** Sequencing reads with pattern 6 repeats that map to chr 1: 11,637,827 do not agree with the assembled sequence. Instead of having parts of the ~35 kb sequence from 11,637,828 bp to the end of the assembly, the reads have a ~1490-bp subtelomeric sequence followed by tandem repeats of pattern 6. **c,** Sequencing reads with pattern 6 repeats support the corrected chr 1 end with the ~1490-bp sequence and pattern 6 tandem repeats.
